# A Versatile Functional Interaction between Electrically Silent K_V_ Subunits and K_V_7 Potassium Channels

**DOI:** 10.1101/2024.02.23.581732

**Authors:** Vijay Renigunta, Nermina Xhaferri, Imran Gousebasha Shaikh, Jonathan Schlegel, Rajeshwari Bisen, Ilaria Sanvido, Theodora Kalpachidou, Kai Kummer, Dominik Oliver, Michael G. Leitner, Moritz Lindner

## Abstract

Voltage-gated K^+^ (K_V_) channels govern K+-ion flux across cell membranes in response to changes in membrane potential. They are formed by the assembly of four subunits, typically from the same family. Electrically silent K_V_ channels (K_V_S), however, are unable to conduct currents on their own. It has been assumed that these K_V_S must obligatorily assemble with subunits from the K_V_2 family into heterotetrameric channels, thereby giving raise to currents distinct from those of homomeric K_V_2 channels.

Herein, we show that K_V_S subunits indeed also modulate the activity, biophysical properties and surface expression of recombinant K_V_7 isoforms in a subunit-specific manner. Employing co-immunoprecipitation, and proximity labelling, we unveil the spatial coexistence of K_V_S and K_V_7 within a single protein complex. Electrophysiological experiments further indicate functional interaction and probably heterotetramer formation. Finally, single-cell transcriptomic analyses identify native cell types in which this K_V_S and K_V_7 interaction may occur. Our finding demonstrate that K_V_ cross-family interaction is much more versatile than previously thought – possibly serving nature to shape potassium conductance to the needs of individual cell types.

## Introduction

Voltage-gated potassium channels (K_V_ channels) are a diverse family of evolutionarily conserved membrane proteins that allow the flux of K^+^ ions across cellular membranes. They play a significant role in determining the excitability of tissues including heart, skeletal muscle, brain, and retina. Based on sequence homology, mammalian K_V_ channels are divided into 12 subfamilies (K_V_1–12) (Alexander *et al*, 2017). Functional K_V_ channels are formed as tetramers of pore-forming α-subunits. These may be made-up by four identical isoforms (homotetramers) or of α-subunits from distinct members of the same K_V_ subfamily (heterotetramers). Heterotetramerization across family borders is typically not possible. The only known exception to this is the heterotetramerization of K_V_2 channels with members of the so-called silent modifier (K_V_S) families, K_V_5, K_V_6, K_V_8, and K_V_9.

K_V_S were termed “silent”, as they are unable to form functional homotetramers at the plasma membrane on their own. When expressed alone, they are retained in intracellular compartments, without giving rise to an electrical current (Bocksteins, 2016). However, when co-expressed with K_V_2 subunits, they co-assemble into functional K_V_S/K_V_2 heterotetramers, with electrophysiological and pharmacological characteristics markedly different to those of K_V_2 channels alone (Bocksteins, 2016; Bocksteins & Snyders, 2012). Such K_V_2-K_V_S heteromerization greatly broadens the functional diversity of K_V_2 channels. The physiological relevance thereof is highlighted by the fact that mutations in the *KCNV2* gene, encoding K_V_8.2 subunits, cause *KCNV2*-associated retinopathy (Gayet-Primo *et al*, 2018; Guimaraes *et al*, 2020). In this condition, the lack of intact K_V_8.2 causes alterations in a conductance termed I_K,x_, which counterbalances the dark current in photoreceptors, to an extent that finally photoreceptor death is caused (Gayet-Primo *et al*., 2018; Hart *et al*, 2019; Smith *et al*, 2012). Similarly, variants in *KCNV1* (K_V_8.1) and *KCNV2* are associated with certain forms of epilepsy (Bergren *et al*, 2009; Jorge *et al*, 2011; Winden *et al*, 2015).

Structurally, the K_V_ α-subunit consists of six transmembrane domains (S1–S6) and a pore loop (between S5 and S6) with the signature sequence (TTIGYGD) determining K^+^ selectivity. The S1–S4 region represents the voltage-sensing domain, and the cytoplasmic N- and C-termini are relevant for subunit assembly, trafficking and functional channel regulation through signalling cascades. Notably, it is well established that K_V_S affect Kv2 currents by altering membrane trafficking, but also by modifying the voltage dependence of activation and inactivation as well as the gating properties (Bocksteins, 2016; Bocksteins & Snyders, 2012; Stas *et al*, 2015). Tetramerization in most K_V_ channels, including the interaction between K_V_2 and K_V_S, primarily occurs in the endoplasmic reticulum (ER). This process involves the primary interaction of T1 domains located in the cytoplasmic N terminus of each subunit (Tu *et al*, 1996). The exact mechanism by which the T1 domain determines subunit specificity remains poorly understood. It is worth noting that certain K_V_ subunits, such as the K_V_7 channel families, lack the T1 domain but are still capable of forming functional homo- and heteromers (Bal *et al*, 2008; Haitin & Attali, 2008; Howard *et al*, 2007). Given the specific co-assembly of closely-related family members, it is reasonable to assume that sequence homology is an important determinant of tetramerization. In this regard, we found that K_V_S (with the exception of K_V_5) are more closely related to members of the K_V_7 family than to the other K_V_ subunits including K_V_2 ((Gutman *et al*, 2005) and Supplementary Figure 1). We therefore hypothesised that K_V_S may constitute modifiers of K_V_7 channels and investigated this interaction in a series of electrophysiological, molecular and cell biological experiments. We found that K_V_S modulate current amplitude and voltage-dependence of neuronal K_V_7 isoforms in a bi-directional manner, and we present biochemical and electrophysiological evidence that K_V_7 and K_V_S physically assemble into the same protein complex, allowing for direct interaction and possibly even heterotetramerisation.

Overall, this work reveals previously unknown ability of K_V_7 and K_V_S to assemble into complexes with unique electrophysiological properties. This may represent a mechanism to extend the native repertoire of K^+^ currents to maintain cell physiology in various tissues.

## Results

### Bi-directional modulation of K_V_7 channels through K_V_S

To explore the potential modulation of K_V_7-mediated currents by K_V_S, we initiated our analyses by characterising K^+^ currents in Chinese hamster ovary (CHO) cells transiently co-expressing K_V_7.2 channel subunits together with different K_V_S in whole cell patch-clamp recordings. Depolarising voltage steps (−100 mV to +60 mV) induced slowly activating outwardly rectifying K^+^ currents that were completely absent in non-transfected CHO cells and in cells expressing only K_V_S (not shown). Strikingly, voltage-dependent currents were significantly reduced in cells co-expressing K_V_7.2 with either K_V_5.1 K_V_8.2, K_V_9.1 or K_V_9.2 subunits compared to cells expressing K_V_7.2 alone (Figure 1 A, B; exemplary recordings are shown in Figure 5 B). Also, in cells co-expressing K_V_7.2 with either K_V_5.1, K_V_8.2, K_V_9.1 or K_V_9.2 we observed an alteration of biophysical characteristics in that the steady-state voltage-dependence of whole cell currents were shifted to significantly more depolarised values as compared to cells expressing K_V_7.2 only. The magnitude of this shift was in the range of about +10 mV for V_h_ values (Figure 1 C, D). The voltage sensitivity as determined by the slope of the voltage dependence was not altered by co-expression of K_V_S. The smaller current amplitudes and altered voltage-dependence were accompanied by significantly more depolarised resting potentials in the cells co-expressing K_V_7.2 with either K_V_5.1 K_V_8.2, K_V_9.1 or K_V_9.2 (not shown). In contrast, steady-state current amplitudes were significantly increased in cells co-expressing K_V_6.1 and K_V_8.1 channels without however affecting voltage dependence or membrane potentials compared to cells expressing only K_V_7.2 subunits (Figure 1 A-D). Co-expression of K_V_6.3, K_V_6.4 or K_V_9.3 did not affect the characteristics of currents through K_V_7.2 channels (Figure 1 A-D).

**Figure 1:**
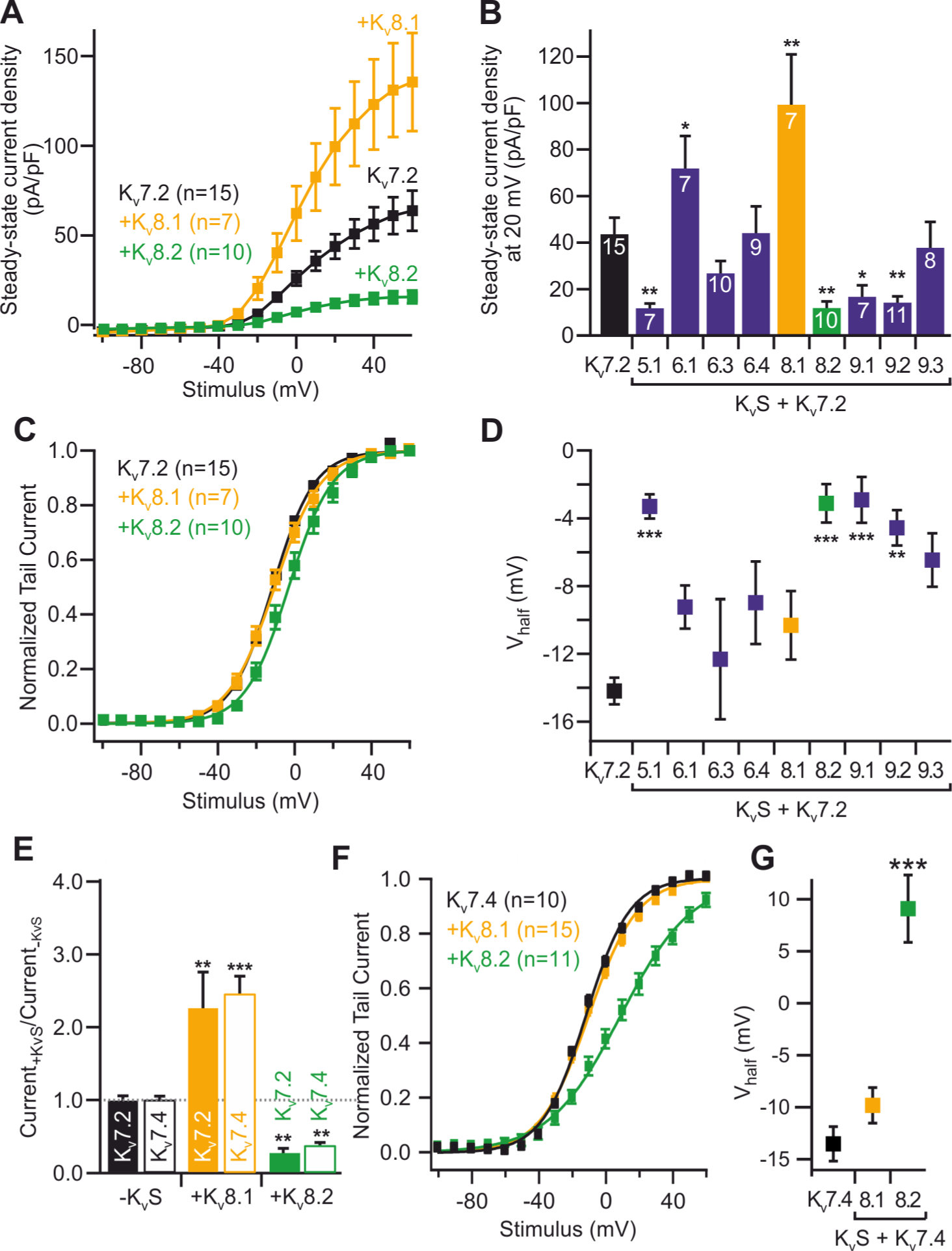
Electrophysiological properties of CHO cells expressing K_V_7 channels are altered by co-expression of K_V_S. **(A)** Voltage-dependent current densities in cells expressing K_v_7.2 alone (black), and together with K_V_8.1 (orange) or K_V_8.2 (green). **(B)** Summary statistics for steady-state current densities at +20mV obtained in recordings similar to those shown in (A) for cells co-expressing K_v_7.2 together with the indicated K_V_S. **(C)** Normalized tail currents and half-maximal activation voltage **(D)** as deduced from Boltzmann fits (solid lines in C) for the same cells as show in A and B. **(E)** Steady-state current densities at +20mV for K_V_7.2 (solid bars) or K_V_7.4 (empty bars) co-transfected with the indicated K_V_S. Data are shown as relative current densities normalized to the mean of current densities observed when K_v_7.2 or K_v_7.4 were transfected alone (“-K_V_S”). **(F)** Normalized tail currents and **(G)** half-maximal activation voltage as deduced from Boltzmann fits (solid lines in F) for cells expressing K_V_7.4 alone (black) or together with the indicated K_V_S.

To assess whether K_V_S similarly modulated the properties of currents through other K_V_7 channels, we performed analogous experiments in CHO cells expressing either K_V_7.4 subunits alone or together with K_V_S. Notably, the effects of co-expression of K_V_S on K_V_7.4 subunits were very similar to those observed for K_V_7.2 channels, except that no effect was observed for K_V_5.1 and shifts in V_h_ were substantially more pronounced (Figure 1 E-G and Supplementary Figure 2).

Taken together, these experiments demonstrated that K_V_S modulated K_V_7 channels in a subunit-specific manner.

### K_V_8 and K_V_7 subunits exist in multi-protein complexes in living cells

We then turned our attention to investigating how K_V_S subunits modulates K_V_7 channels at the molecular level, focusing on K_V_8.1 and K_V_8.2 subunits and analysed whether K_V_7 and K_V_8 subunits were present in the same protein complex and in close proximity to each other. We hence carried-out BioID and co-immunoprecipitation experiments in HEK cells. For BioID experiments, we transiently overexpressed BioID2-HA-tagged K_V_S (K_V_8.1 or K_V_8.2) and flag-tagged K_V_7 subunits (Figure 2). To ensure that the BioID2 fusion did not affect the expression and subcellular localisation of the K_V_S subunits, we performed immunostainings on transfected HeLa cells with antibodies against the HA tag at the C-terminal end of K_V_8.1, K_V_8.2 BioID2 fusion proteins. This revealed the expected intracellular localisation with a signal pattern typical of the endoplasmic reticulum (ER) within the cells. (Figure 2 B; Supplementary Figure 8).

**Figure 2:**
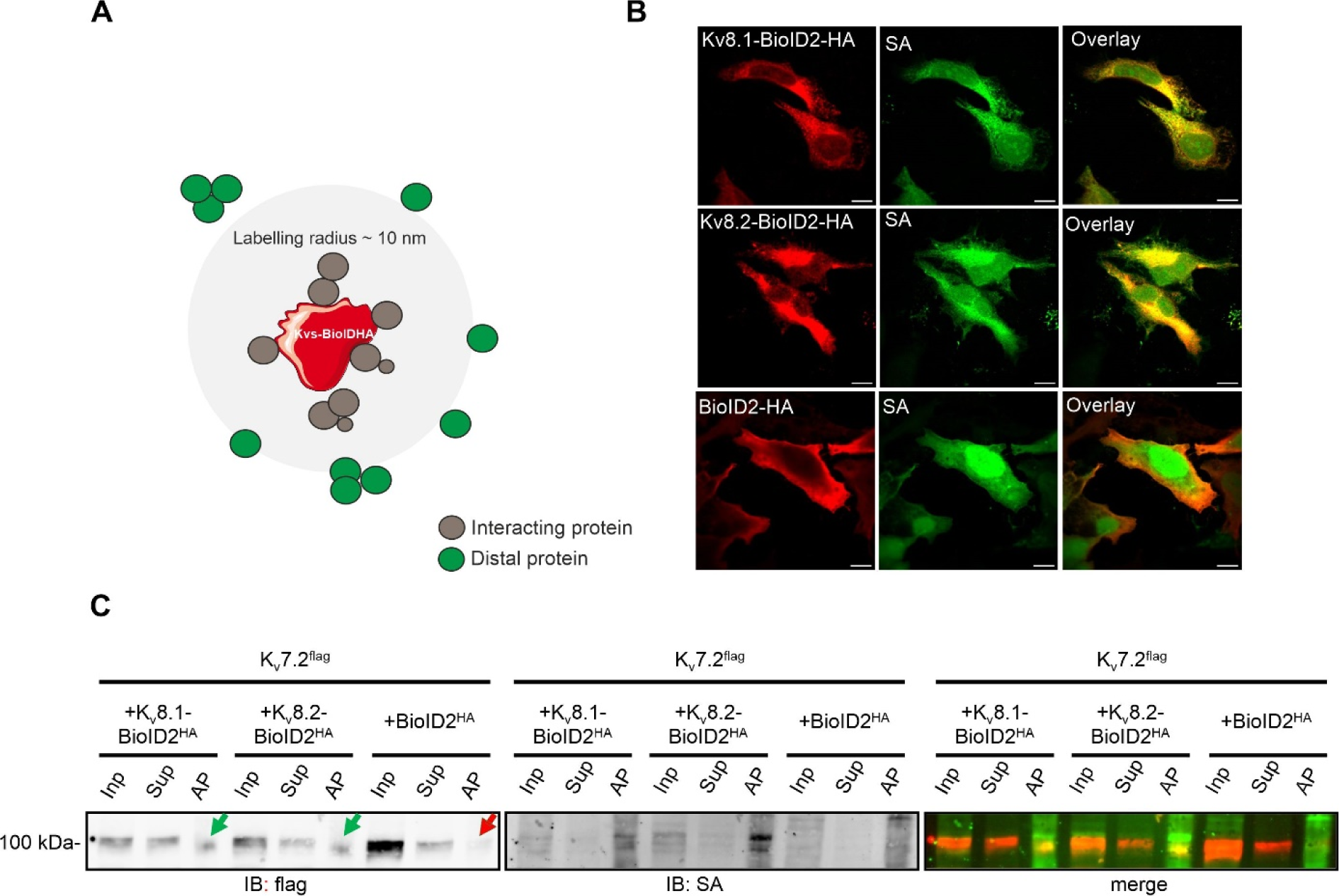
K_V_7 and K_V_S assemble in close proximity. **(A)** BioID assay methodology. The K_V_S (8.1 / 8.2) was fused to the promiscuous form (BirA*) of the bacterial biotin ligase BirA and co-expressed with flag tagged K_V_7.2 subunits in HEK293 cells. Upon the addition of biotin, proximal proteins (grey) were biotinylated within a labeling radius of ∼10 nm, whereas the distal proteins (green), remained unlabelled. **(B)** Representative confocal images of HEK293 cells transiently transfected with BioID2-HA tagged K_V_8.1 (first row), BioID2-HA tagged Kv8.2 (second row), BioID2-HA alone (third row). Cells were stained with either HA antibody (red, first column) or Alexa Fluor™ 488 streptavidin conjugate (green, second column). Scale bar, 10 μm. **(C)** Following biotin labelling, cells were lysed and biotinylated proteins were then purified using streptavidin beads and identified by western blot analysis. Immunoblotting (IB) was performed by using an anti-flag antibody to detect the flag-labelled Kv7.2 subunits and an Alexa Fluor™ 488 streptavidin conjugate (SA) to detect the biotinylated proteins. Note the absence of flag tagged Kv7.2 subunits in the affinity purified (AP) lane with BioID2-HA alone (indicated by red arrow). However, they are present in both the (AP) lanes (indicated by green arrows). Overlaying the signals obtained from anti-flag and streptavidin staining revealed a noticeable proximity of these distinct bands (merge). Similar results were obtained in n = 3 transfections.

Biotinylation of endogenous proteins in cells expressing HA-BioID2-KvS (8.1 and 8.2) with exogenous biotin strongly stimulated a wide range of endogenous proteins on western blots probed with Alexa Fluor™ 488 streptavidin conjugate (data not shown). This indicates that the BioID2 moiety was adequately exposed in the K_V_S fusion construct, allowing for efficient biotinylation. Our next step was to test whether the Kv7 subunits are in close proximity to the K_V_S subunits. If they are indeed in close proximity, they should be biotinylated and then precipitated with streptavidin beads. To achieve this, we transiently expressed BioID2 tagged K_V_S (8.1 or 8.2) and flag tagged K_V_7.2 constructs in HEK293 cells in the presence of 50 µM biotin for 24 h and then lysed the cells using a radio-immunoprecipitation assay (RIPA) lysis buffer. HEK293 cells transfected with BioID2-HA alone, processed in parallel, were used as negative controls. As shown in Figure 2C (anti-flag staining, green arrows), the flag-tagged K_V_7.2 subunits were robustly precipitated using streptavidin beads, strongly suggesting that the K_V_7.2 subunits are indeed in close proximity (10 nm distance) to the K_V_S (8.1 and 8.2) subunits and are therefore biotinylated, whereas no precipitation occurred when BioID2 alone was used as a negative control (red arrow). To complement these data with an independent approach, we performed a proximity ligation assay (PLA) on cells transfected with myc-tagged K_V_S and flag-tagged K_V_7. Indeed, PLA signals were only observed in cells co-expressing both, K_V_S and K_V_7 (Supplementary Figure 3).

We then asked whether the flag-tagged K_V_7.2 was located in the same complex as the K_V_S subunits by carrying out co-immunoprecipitation experiments. The protein complexes were precipitated with anti-flag M2 beads, and the subunits in the precipitates were detected in western blots with antibodies directed against myc or flag tags, respectively (Figure 3 A, B). These results demonstrated that the K_V_7.2 and K_V_S channels were not only located in close proximity, but also exist in a single multi-protein complex. As a negative control, co-immunoprecipitation experiments were carried out using Dynabeads coated with unrelated IgG antibodies and with cell lysates devoid of KCNQ subunits (Supplementary Figure 4, Supplementary Figure 5), demonstrating the specificity of co-immunoprecipitations.

**Figure 3:**
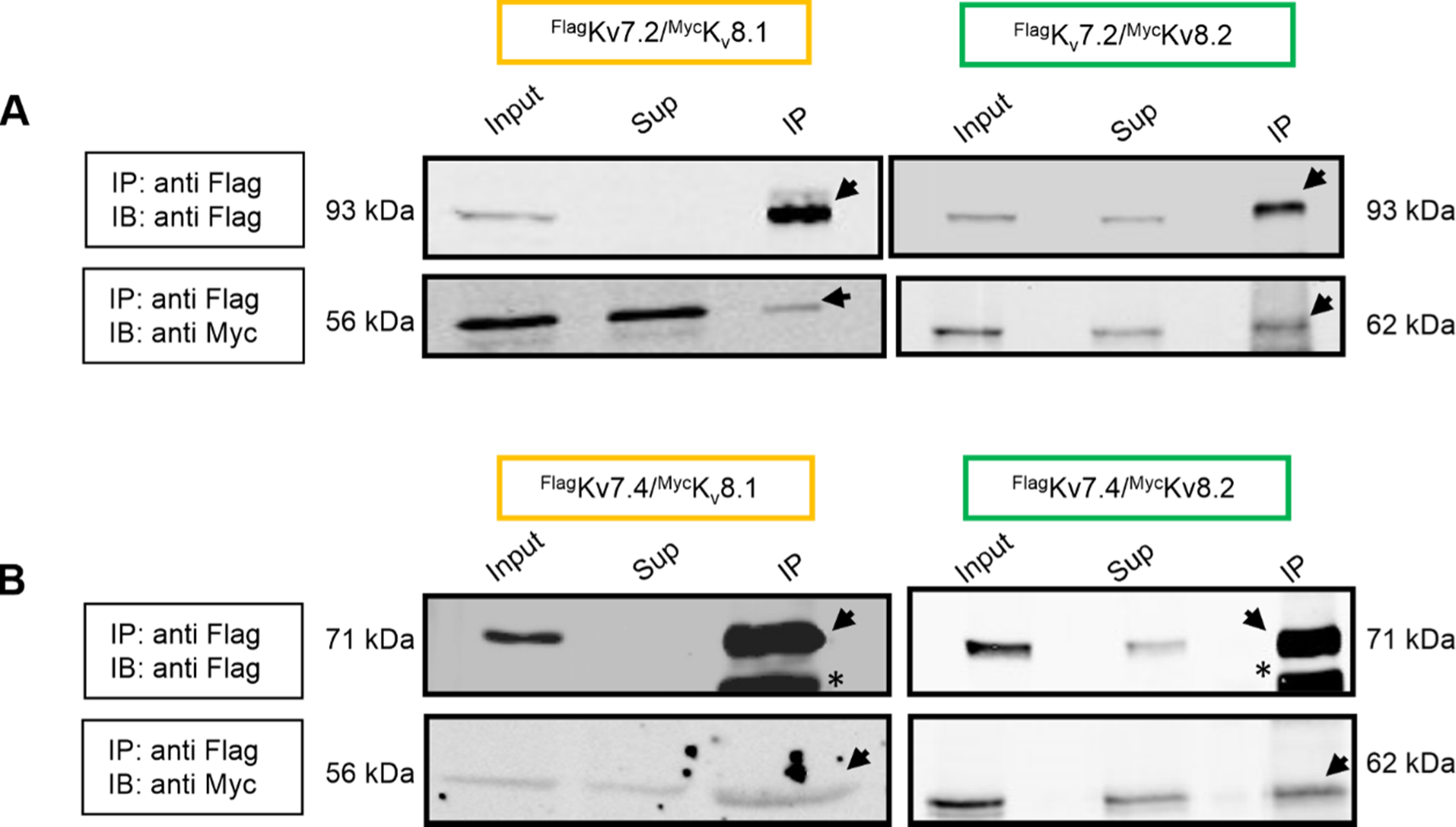
Kv7 and KvS assemble into a protein complex. Co-immunoprecipitation experiments were performed using myc-tagged Kv8.1 (left) and Kv8.2 (right) subunits in combination with flag-tagged Kv7.2 (A) and Kv7.4 (B) channel subunits from HEK293 cell lysates using monoclonal anti-flag M2 conjugated agarose beads. The precipitate was then subjected to western blot analysis using monoclonal anti-flag M2 flag antibody and anti-myc antibodies respectively. Note the presence of both K_V_7 and K_V_S subunits in the precipitated fraction (indicated by black arrows). Asterisks indicate heavy chains detected by secondary antibodies. Abbreviations used: IP (immunoprecipitation), Sup (supernatant), IB (immunoblotting). Similar results were obtained in n = 3 transfections.

### K_V_8 modulate K_V_7 currents by affecting both membrane trafficking and biophysical properties

We then sought to determine whether K_V_S modulated K_V_7 currents through mechanisms similar to those involved in the regulation of K_V_2 channels. To this end, we measured the plasma membrane expression of both K_V_7.2 and K_V_7.4 channels containing an extracellular HA tag, in *Xenopus laevis* oocytes using a luminometric assay (Renigunta *et al*, 2014). This method involves the oxidation of luminol by horseradish peroxidase (HRP) in conjunction with antibodies that can detect HA-tagged K_V_7 channels located on the oocyte surface. The intensity of light emitted directly correlates with the membrane expression of K_V_7 channels (Figure 4 A). In these experiments, co-expression of K_V_8.1 significantly increased the membrane expression of both K_V_7.2 and K_V_7.4 channels, whereas co-expression of K_V_8.2 slightly decreased the membrane expression of both channels (Figure 4 B, C), fully in line with the observed effects on current density (c.f., Figure 1). Taken together, these data indicated that K_V_8.1 enhances K_V_7 currents by increasing their membrane expression, whereas co-expression of K_V_8.2 may reduce current amplitudes, at least in part, by attenuating surface expression of K_V_7 channels. Therefore, we conclude that K_V_S modulates K_V_7 currents by altering their membrane abundance, similar to the mechanisms reported for K_V_2.1 channels, but, unlike K_V_2 channels, in a bidirectional manner (Bocksteins, 2016; Bocksteins & Snyders, 2012).

**Figure 4:**
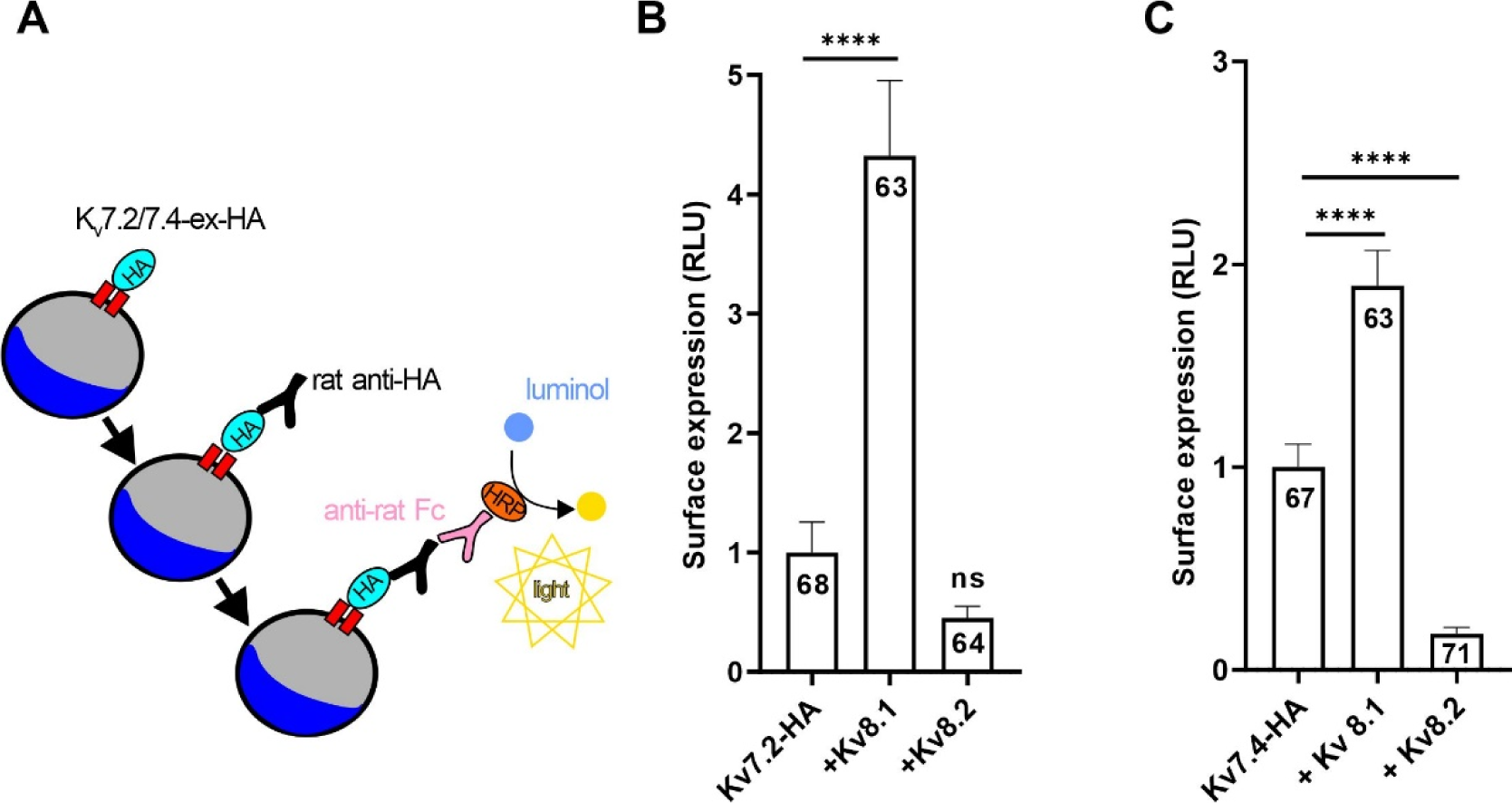
K_v_7 and K_v_S form heteromers with altered membrane trafficking. **(A)** Surface expression of HA-tagged K_v_7 channels measured with a luminometric technique in Xenopus oocytes. Mean surface expression of **(B**) HA-tagged K_v_7.2 channels **(C**) HA-tagged K_v_7.4 channels (measured in relative light units [RLUs]) in Xenopus oocytes after injection of HA-tagged K_v_7 cRNA either alone or together with K_v_8.1 or K_v_8.2. Uninjected oocytes were used as negative control.

We then set out to gain insight into whether K_V_S subunits modulate K_V_7 channels via heterotetramerization into the same channel complex or by close interaction, e.g., as associated but independent channel entities or as some sort of ancillary beta subunit (Figure 5 A). To determine whether all isoforms contribute to the channel pore (i.e. heterotetramerization), we generated channels containing mutations in the GYG pore motif (GYG/AAA exchange for K_V_8.1 and GYG/GYD exchange for K_V_7.2) that render these variants inactive. When co-expressed with wild-type subunits, such variants are known to attenuate whole cell currents via dominant-negative effects (Leitner *et al*, 2012). Assuming fully stochastic co-assembly of equally available subunits, co-expression of wild-type and mutant subunits is predicted to reduce whole cell current amplitudes to 1/16 (compared to cells expressing only wild-type subunits), leaving intact a minimal number of channels containing only wild-type subunits (illustrated in Figure 5 A, first row). We performed patch-clamp recordings on cells expressing wild-type and mutant isoforms taking current densities as measure for potential co-assembly of the subunits. Co-expression of wild-type K_V_8.1 with wild-type K_V_7.2 significantly increased whole cell current amplitudes compared to cells expressing only K_V_7.2 channels, as we had observed before (Figure 5 B). When we co-expressed pore-mutated K_V_7.2(GYD) with wild-type K_V_8.1, current amplitudes were reduced to virtually zero. These experiments demonstrated that K_V_8.1 subunits alone were not able to form an independent functional pore (Figure 5 B).

**Figure 5:**
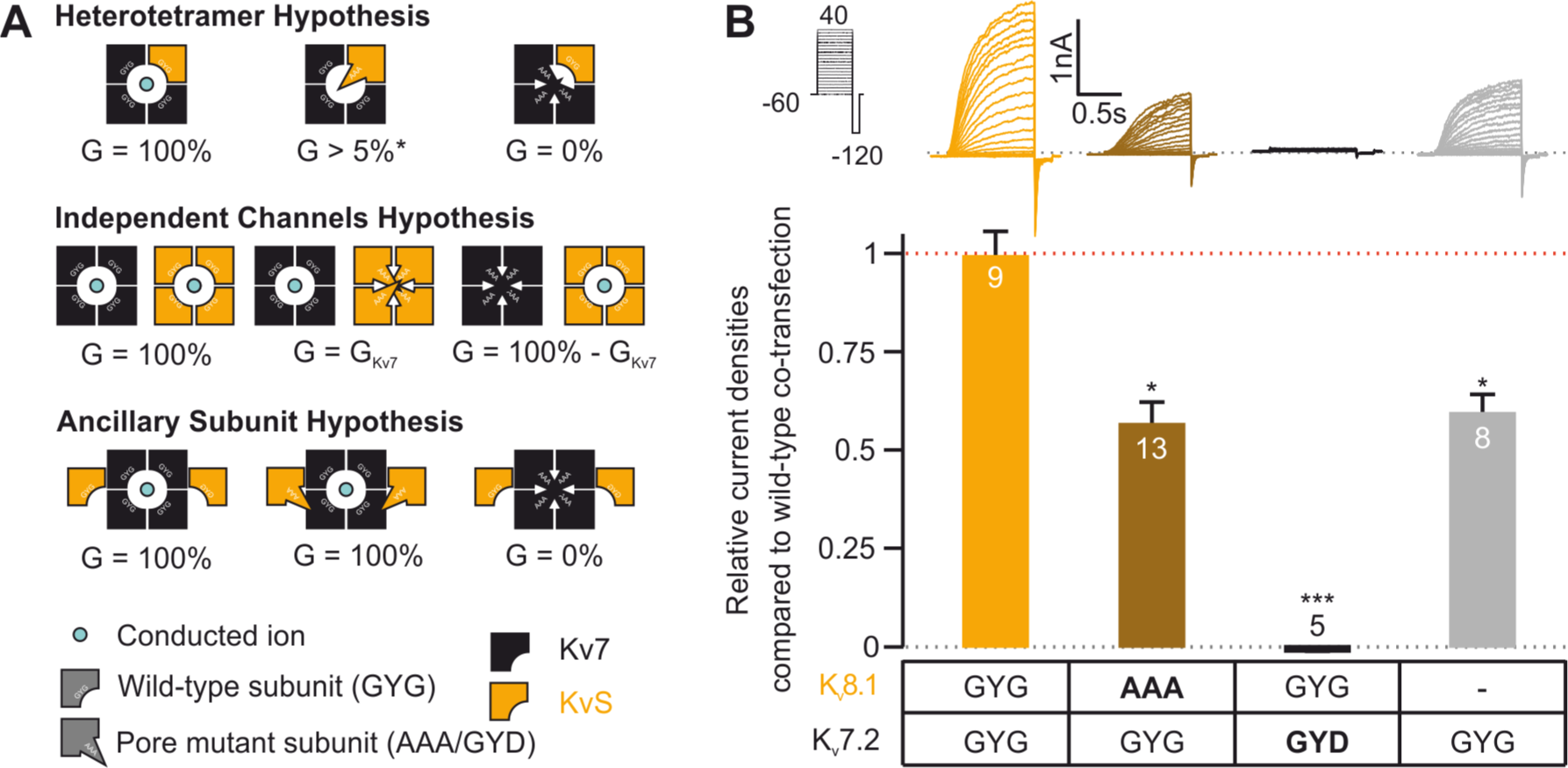
K_v_7 and K_v_S form heteromers with a single conducting pore. **(A)** Imaginable modes of interaction of K_V_7 with K_V_S, consequences of expression of pore mutant subunits and expected membrane conductance. *) The percentage of current reduction depends on the probability of inclusion of the pore-mutant subunit into a heterotetramer. Detailed explanation is given in the main text. **(B)** Patch-clamp recordings from cells co-transfected with either combination of wild-type (GYG) and pore-mutated (AAA, GYD, resp.) K_V_8.1 and K_V_7.2 or with K_V_7.2 alone. Top panel shows stimulus protocol (black) and exemplary recordings. Bottom panel shows summary statistics for steady-state current densities at +20mV. Data are shown as relative current densities normalized to the mean of current densities observed when transfecting wild-type K_V_7.2 with wild type K_V_8.1.

When wild type K_V_7.2 was transfected together with pore-mutated K_V_8.1(AAA), current densities were reduced to approximately 50% of those observed when measuring cells transfected with both wild type constructs, resulting in current densities minimally smaller than those observed in cells transfected with K_V_7.2 alone (Figure 5 B, note that amount of cDNA encoding K_V_7.2 was kept constant). These results suggested that functional pore regions of K_V_8.1 are essential for K_V_S-dependent modulation of K_V_7 channels. Noteworthy, the (only) 50% current reduction observed upon K_V_8.1(AAA) co-transfection may indicate that during co-assembly K_V_7.2 homotetramer formation is more likely to occur than heterotetramer formation. In particular, the 50% reduction observed herein would be expected if the probability for K_V_7.2 homomers was approximately 5-fold higher than that for K_V_7.2-K_V_8.1 heteromers. Yet, our observations clearly rule out alternative modes of interaction: If K_V_7.2 and K_V_8.1 co-existed as independent channels (Figure 5 A, second row), a substantial current should have been observable upon co-expression of pore-mutated K_V_7.2(GYD); If K_V_8.1was a beta-subunit to K_V_7.2 (Figure 5 A, third row), mutating the “pore” sequence of K_V_8.1 would not be expected to have any effect at all. Taken together, these data suggested that K_V_7 and K_V_8.1 subunits can likely heteromerise into functional channels with a slight preference for K_V_7 homomers.

### K_V_7 and K_V_S are expressed in the same organs and cells

To investigate whether an interaction between K_V_7 and K_V_S might be physiologically relevant, we used reverse transcription quantitative polymerase chain reaction (RT-qPCR) to examine the expression of K_V_8 in three tissues, where the role of K_V_7 channels is well established (Wang & Li, 2016). Indeed, we found that in hippocampus and dorsal root ganglia, the abundance of *Kcnv1* (K_V_8.1) mRNA was in the same order of magnitude as that of *Kcnq2* (K_V_7.2) mRNA. In both these neural tissues, also *Kcnv2* (K_V_8.2) mRNA was detected, though at much lower levels (Figure 6 A). By contrast, in heart, expression levels of *Kcnv2* (K_V_8.2) were comparable to those of *Kcnq1* (K_V_7.1), while *Kcnv1* mRNA was not detected (Figure 6 B).

**Figure 6:**
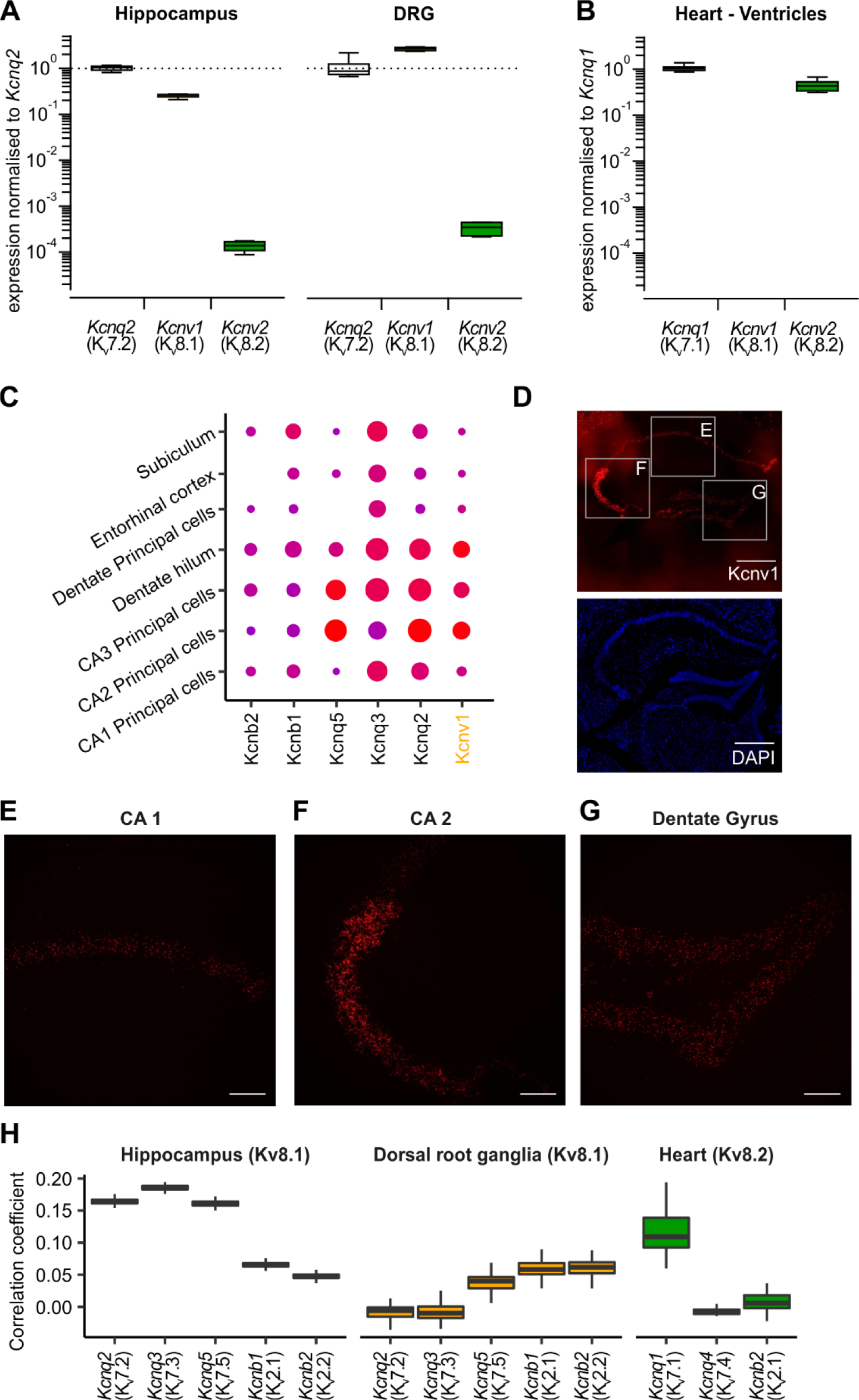
K_V_7 and K_V_S co-express in various tissues in a cell specific manner. **(A, B)** Reverse transcription quantitative PCR for K_V_7 and K_v_S transcripts from neuronal (A) and cardiac (B) tissue. For each condition 8 biological replicates with two technical replicates were included into the analysis. Shown are expression levels normalized to the mean expression level of K_V_7.2 for Hippocampus and Dorsal Root Ganglia (DRG) (A) and K_V_7.1 for Heart (B), respectively. **(C)** Expression dot plots of K_V_8.1, K_V_7 and K_V_2 genes, for comparison, in Hippocampus. Shown are only clusters with K_v_8.1 expression. The complete dot plot is given in Supplementary Figure 7. **(D-G)** RNAScope of K_V_8.1 in the mouse hippocampus. **(D)** Low-magnification overview. Scale bar: 500 μm. **(E-G)** High-magnification confocal micrographs of CA 1 (E), CA 2 (F) regions and Dentate Gyrus (G), respectively. Scale bar: 100 μm. **(H)** Correlation coefficients for the single-cell expression of K_V_7 and K_V_2 with K_V_8.1 or K_V_8.2, respectively. Shown are Pearson’s correlation coefficients for log2+1-transformed transcript counts from individual cells as observed in three publicly available singe-cell RNA sequencing datasets. Included into the analysis were only cell types found to express the respective K_V_S.

To further evaluate whether K_V_7 and K_V_S transcripts merely coexist in the same tissue, or are actually transcribed in the same cells we analysed publicly available single-cell RNA sequencing (scRNAseq) datasets from the hippocampus (Saunders *et al*, 2018), dorsal root ganglia (DRG) (Finno *et al*, 2019) and the heart (Tabula Muris *et al*, 2018). Consistent with our observations in the qPCR experiments we found high read counts for *Kcnq2* and *Kcnv1* in hippocampal and DRG cells, whereas *Kcnv2* was barely detected. In the dataset from cardiac cells, in turn, *Kcnq1* and *Kcnv2* were predominant, whereas Kv8.1/*Kcnv1* was not found (Supplement material file 1, Table S2).

In the hippocampus, *Kcnv1* was encountered in seven clusters, with the highest transcript counts being observed in the clusters representing CA1-3 pyramidal neurons, dentate hilus and dentate principal neurons (Figure 6 C). To see if the observed transcriptomic expression pattern would also be supported by methods that provide spatially encoded information, we employed single molecule fluorescence RNA in-situ hybridization (“RNAScope”). Indeed, using RNAScope we find *Kcnv1* expression patterns closely matching the observations made using scRNAseq (Figure 6 D-G). By far the highest *Kcnv1* signal was obtained from the CA2 region, followed by the dentate gyrus and the CA1 region. In scRNAseq, both K_V_7.2/*Kcnq2* and K_V_2.1/*Kcnb1* were both found in each of these cell types /clusters (Figure 6 H, Supplementary Figure 7 A). We argued that if K_V_8.1 would interact with K_V_7.2 rather than a K_V_2 in hippocampal neurons, one would expect *Kcnv1* expression levels to correlate stronger with those of *Kcnq2* than with those of *Kcnb1* or *Kcnb2* at the single cell level. Indeed, Pearson’s correlation coefficient (PCC) for *Kcnv1* with *Kcnq2* was higher than the correlation of *Kcnv1* with 99 [98 - 99] % of all other transcripts encountered, and in particular higher than that with *Kcnb1* (Figure 6 H, left, Supplement material file 1, Table S3).

In dorsal root ganglia, Kcnv1 and *Kcnq2* overlapped in two clusters (Supplementary Figure 7 B). In stark contrast to the hippocampus, *Kcnv1* expression levels in DRG were more strongly correlated with those of K_V_2 channel genes rather than with those of K_V_7 channel genes (Figure 6 H, middle, Supplement material file 1, Table S3). In the heart, *Kcnv2* was found exclusively in the cluster representing cardiac myocytes, as was *Kcnq1* (Supplementary Figure 7 C). Here, the PCC for *Kcnv2* with *Kcnq1* was higher than the correlation of Kcnv2 with 0.99 [0.98 - 1.00] % of all transcripts encountered, and, again, particularly higher than that with Kcnb1 (Figure 6 H, right, Supplement material file 1, Table S3). Taken together, these transcriptomic analyses suggest that K_V_7 and K_V_8 mRNA expression is strongly correlated within individual cells in hippocampus and heart, whereas this was not the case in the DRG.

## Discussion

In the present work, we show that K_V_ channels from the silent K_V_-subfamily (K_V_S) can interact with K_V_7 channels to alter their trafficking and biophysical properties. Modulation of K_V_2 channels through K_V_S has long been reported (Bocksteins, 2016; Bocksteins & Snyders, 2012) and, up until now, this has been considered to be the only case of cross-family interaction among K_V_ channel families. Yet, in analogy to their actions on K_V_2 channels (Bocksteins, 2016), K_V_S reduced K_V_7-mediated currents through an alteration of K_V_7 membrane targeting and voltage-dependency. Importantly, – and in contrast to their effects on K_V_2 – certain K_V_S also increased K_V_7 current amplitudes and surface expression. Thus, modulation of K_V_7 by K_V_S is bidirectional and thereby more diverse than the modulation of K_V_2. Based on our data, we hypothesise that K_V_7 co-assemble with K_V_S into functional heteromers. This further broadens the functional diversity of K_V_ channels and may serve to shape potassium conductance to fit the needs of individual cell types.

### Mode and functional aspects of the K_V_7-K_V_S interaction

Using PLA and BioID experiments, we herein demonstrated that in living cells K_V_7 co-exist with K_V_8.1 and K_V_8.2 in close physical proximity. In addition, we could show by co-immunoprecipitation that these ion channels assemble into a common protein complex. This interaction dictates trafficking of K_V_7 channels to the plasma membrane allowing for a bidirectional modulation of K_V_7-mediated current amplitudes. Indeed, the enhanced surface abundance of K_V_7.2 and K_V_7.4 in *Xenopus laevis* oocytes upon co-expression of K_V_8.1 was paralleled by increased macroscopic K_V_7.2 and K_V_7.4 current amplitudes recorded from CHO cells. Conversely, co-expression of K_V_8.2 resulted in a reduction in both macroscopic current amplitudes and surface expression. We conclude that the molecular mechanism of the K_V_S-dependent modulation of K_V_7 currents may be similar to that of K_V_2 channels (i.e., modulation of surface expression plus modulation of biophysical properties). However, bidirectional modulation of current amplitudes indicates that the interaction between K_V_7 and K_V_S may be somewhat more versatile than their interaction with K_V_2 channels, which are generally down-regulated by K_V_S (Bocksteins, 2016). Taking into account our RNA-sequencing analyses, co-expression of K_v_7.2 and K_v_8.1 in hippocampal neurons, for example, may lead to a tuned decrease in electrical excitability by increasing the Kv7-medited M-current (Wang *et al*, 1998). Modulation of K_V_7.1 by K_V_8.2 subunits in cardiac muscle, in turn, may lead to a slower and delayed repolarization process. However, mouse models carrying genetic deletions will be required to unequivocally demonstrate whether this novel interaction is also relevant in native tissues.

Although our data provide clear evidence for the coexistence of K_V_7 and K_V_S subunits in a common protein complex, and the functional relevance of the interaction, we are unable to draw a definite conclusion as to whether K_V_7 and K_V_S co-assemble into functional heterotetramers, as it has been previously been shown for K_V_2 channels (Bocksteins *et al*, 2009; Moller *et al*, 2020). Since K_V_S altered voltage dependence of K_V_7 channels, we propose heteromerisation also for K_V_S and K_V_7 channels. In support of this notion, we observed that co-transfection of dominant-negative pore-mutant K_V_8 with wild-type K_V_7.2 led to a robust (∼50%) reduction of current amplitudes as compared to transfection of both wild-type subunits, i.e., the functional modulation of K_V_7.2 currents was abolished via co-expression of K_V_8 pore mutants. While these findings are in line with a possible heterotetramerization, the dominant-negative effects of K_V_S pore mutants on K_V_7 currents were somewhat weaker than expected: Assuming each subunit would be built into a heterotetramer with equal probability a ∼95 % reduction in current amplitude would be anticipated. Importantly, it is unclear whether the potential K_V_7-K_V_S heterotetramerization is fully stochastic. Specifically, a roughly 5-fold assembly preference for respective homotetramers over heterotetramers would explain the observed 50% current reduction. Potential preference for homomers over heteromer formation might seem surprising, but it has indeed been demonstrated even for the well-studied K_V_2-K_V_8.2 interaction, and the degree of current reduction observed for K_V_2.1 by dominant-negative K_V_8.2 mutants is only slightly higher than what we observed in our study (Czirjak *et al*, 2007; Smith *et al*., 2012). Nevertheless, there is still room for debate regarding the mode of interaction and studying Kv7.4 channels (where changes in the V_h_ upon co-transfection of dominant-negative K_V_S) or on K_V_7-K_V_S concatamers may help to clarify this issue in future.

### The physiological role of Kv7-KvS interaction

In the present study, we used a wide variety of cell types, including cell lines from primates, rodents and amphibian origin. This substantiates the notion that the interaction between K_V_7 and K_V_S is not restricted to a particular species or cell line, but instead represents a universally observable and evolutionarily stable phenomenon.

To further explore the presence of K_V_7-K_V_S interaction in native systems, we delved into where this interaction might occur. Through RT-qPCR analysis, we detected the expression of K_V_8.1 and K_V_8.2 in three tissues, where K_V_7 play a well-established, prominent role (Greene & Hoshi, 2017; Maljevic *et al*, 2008): the hippocampus, dorsal root ganglia, and the heart (Figure 6 A-C). We then analysed single-cell RNA-sequencing datasets, and observed that K_V_7 and K_V_S were not just expressed in the same tissues, but actually in the same cell types and cells (Supplementary Figure 7). Moreover, we found that on the level of individual cells, in several subtypes of hippocampal neurons (e.g. CA1-CA3) and cardiac muscle cells expression levels of K_V_7 and K_V_S were robustly correlated. Furthermore, our analysis revealed that in the hippocampus and heart, the expression of K_V_8.1 and K_V_8.2, respectively, showed a substantially stronger correlation with K_V_7 as compared to K_V_2. Correlation in transcript counts obviously do not immediately proof functional interaction on protein level. Thus, these findings fall short in providing direct evidence for a functional K_V_7-K_V_S interaction in the examined tissues. Recent studies, however, show that transcript-level correlation is commonly maintained on protein level and can help to predict protein function (Ribeiro *et al*, 2022; Sharan *et al*, 2007).

To establish definitive evidence for the presence of K_V_7-K_V_S complexes in native tissues and unravel their physiological significance, further investigations encompassing both structural and functional studies are warranted. Specifically, functional studies present significant challenges that can only be addressed by establishing knockout mouse models and/or a comprehensive pharmacological characterization of K_V_7/K_V_S complexes. In this regard, it is encouraging that a first K_V_8.2 mouse model is available and has been partially characterized (Hart *et al*., 2019; Jiang *et al*, 2021). These approaches will clarify the specific roles and functional properties of K_V_7-K_V_S interactions in biological systems.

### Translational relevance

While this is primarily a cell physiological study, the herein-described K_V_7-K_V_S interaction may hold promise for clinical and translational applications. The newfound K_V_7-K_V_S complex may pose unique pharmacological properties that are distinct from K_V_7 complexes not encompassing K_V_S. This would enable the development of pharmacological agents specifically targeting K_V_7-K_V_S complexes while leaving K_V_7 homotetramers unaffected. Such an approach could enable highly specific pharmacotherapies e.g. for epilepsy or cardiac arrhythmias.

Indeed, before persuading such translational directions a better understanding of the role of K_V_7-K_V_S complexes in native tissues as well as their pharmacological properties is required.

### Conclusion

In summary, our study provides evidence for the formation of a common protein complex between K_V_S and K_V_7 subunits. These complexes exhibit distinct electrophysiological properties, possibly through heterotetramer formation. Our data, suggest that such interactions may also occur in native tissues, particularly in the hippocampus and the heart. K_V_S-K_V_7 interactions could therefore represent a mechanism utilized by nature to further increase the functional diversity of K_V_ channels and to fine-tune the electrophysiological properties of individual cell types to their functional needs.

## MATERIAL AND METHODS

### Animal studies

Adult female African clawed frogs (*Xenopus laevis*) were used for experiments with Xenopus oocytes. The frogs were anaesthetized by placing them in water containing 1 g/l tricaine. Stage V oocytes were collected from the ovarian lobes. Anesthesia and surgery were performed with the approval of the Giessen Regional Animal Health Authority.

C57Bl/6J mice were purchased from Janvier Labs (Le Genest-Saint-Isle, France). Tissue collection was performed in accordance with the Ethics Guidelines of Animal Care (Medical University of Innsbruck).

### Molecular cloning and mutagenesis

Supplemental Table S1 summarizes the constructs used in this study. QuikChange II XL Site-Directed Mutagenesis Kit (Stratagene, Agilent Technologies, Waldbronn, Germany) was utilized to introduce point mutations. To enhance expression efficiency, all constructs utilized for experiments in Xenopus oocytes were subcloned between the 5’ and 3’ UTR of the Xenopus β-globin gene in the modified pSGEM vector. The mMessage mMachine kit (Ambion, Huntingdon, UK) was used to synthesize complementary RNA transcripts. For surface quantification assays, a plasmid containing KCNQ2 (Schwake *et al*, 2003) or KCNQ4 (HJ Kim *et al*, 2011) with an external hemagglutinin (HA) epitope tag was used. All DNA constructs were verified through Sanger sequencing.

### Cell Culture and Transfection

Chinese hamster ovary (CHO) dhFR^-^ cells were maintained as previously described (Lindner *et al*, 2011). In brief, cells were kept in MEM Alpha Medium supplemented with 10% fetal calf serum (FCS) and 1% penicillin/streptomycin (Invitrogen GmbH, Darmstadt, Germany) in a humidified atmosphere at 5% CO_2_ and 37°C. Cells were transiently transfected with jetPEI transfection reagent (Polyplus Transfection, Illkirch, France). HeLa and HEK cells were cultured in the same way as CHO. Transfection of HeLa cells was performed using jetPRIME transfection reagent (Polyplus Transfection, Illkirch, France).

### Electrophysiological Recordings

Whole-cell recordings were performed on transiently transfected CHO cells in culture, as previously reported (Leitner *et al*, 2011; Leitner *et al*, 2018; Wilke *et al*, 2014). All experiments were performed approximately 48 h after transfection at room temperature (22°C-25°C) (Dierich *et al*, 2020). During recordings, cells were kept in extracellular solution containing (in mM): 144 NaCl, 5.8 KCl, 1.3 CaCl_2_, 0.7 Na_2_HPO_4_, 0.9 MgCl_2_, 5.6 glucose, 10 HEPES, pH adjusted to 7.4 (NaOH) (305-310 mOsm/kg). Whole-cell patch clamp recordings were performed at room temperature (19-23°C) with an HEKA EPC10 USB patch clamp amplifier controlled by PatchMaster software (HEKA, Lambrecht, Germany) or an Axopatch 200B amplifier (Molecular Devices, Union City, CA). Voltage clamp recordings were low-pass filtered at 2.5 kHz and sampled at 5 kHz. Recordings were excluded from analyses, when the series resistance (R_s_) was ≥ 7 MΩ, and R_s_ was compensated through-out the recordings to 80%, with the exception of the data presented in Figure 5, where no R_s_ compensation was performed. Patch pipettes were pulled from borosilicate glass (Sutter Instrument Company, Novato, CA, USA) and had a resistance of 2-3.5 MΩ after filling with intracellular solution containing (mM): 135 KCl, 3.5 MgCl_2_, 2.4 CaCl_2_ (0.1 free Ca^2+^), 5 EGTA, 5 HEPES and 2.5 Na_2_-ATP (pH adjusted with KOH to 7.3; 290-295 mOsm/kg).

### Co-immunoprecipitation

To perform immunoprecipitations, anti-FLAG M2 magnetic beads from Sigma-Aldrich (St. Louis, MO) were used in accordance with the manufacturer’s instructions. Briefly, lysates from HEK293 cells that expressed flag-tagged K_V_***7*** and myc-tagged K_V_S were utilized. The beads were coated with antibody and then incubated with the cell lysates under rotation at 4°C for 12 hours. Beads with antigen-antibody complex were extensively washed (4 times) with wash buffer, followed by elution of the proteins on the beads through boiling at 72°C for 10 minutes in 2 x SDS sample loading buffer. The proteins were then separated on a 10-12% SDS-PAGE under reducing conditions, transferred to a nitrocellulose membrane and probed with either a mouse anti-myc antibody (1:1000; Cell Signaling, Danvers, MA) or a monoclonal ANTI-FLAG® M2 antibody (1:1000; Sigma-Aldrich). Fluorescent secondary antibodies (1:5000; Bio-Rad, Hercules, CA) were used to visualize the membrane, which was then imaged with a ChemiDoc MP imaging system from Bio-Rad.

### Proximity Ligation Assay

CHO cells were fixed in 4% ice-cold methanol-free paraformaldehyde as previously described (Lindner *et al*, 2020). Proximity ligation assay (PLA) was performed using the Duolink *In-Situ* Red Starter Kit (Sigma-Aldrich) according to the manufacturer’s protocol, with the amplification step being performed by incubation with polymerase at 37°C for 50 minutes.

Primary antibodies used were rabbit anti myc (71D10, Cell Signaling) at a dilution of 1:200 and mouse anti flag (F1804, Sigma-Aldrich) at 1:400. Fluorescent secondary antibody (Alexa 488 conjugated Donkey anti mouse, ThermoFisher, Waltham, USA, 1:250) was added during the PLA probe incubation step. Image acquisition was performed using an LSM710 confocal laser scanning microscope (LSM, Carl Zeiss, Jena, Germany) as previously described (Lindner *et al*., 2020).

### BioID Proximity-Dependent Biotinylation assay

K_V_S-BioID2 (HA tag and BirA R118G mutant fused to the C-terminus of K_V_S subunits) was transfected into 6 cm plates of HEK293 cells. After 24 hours of transfection, 50 μM biotin was added to the culture medium to induce biotinylation of proteins in the vicinity of BioID2-K_V_S in the cells for 18-24 hours. Cells were lysed in RIPA lysis buffer (150 mM NaCl; 50 mM Tris HCl pH=7.4; 1% Triton X100; 0.1% SDS; 0.5% sodium deoxycholate). The sample was passed through an 18-gauge needle several times to reduce viscosity. Biotinylated proteins were purified using streptavidin-agarose (Pierce/Thermo Fisher) and eluted in 6x SDS-PAGE sample buffer containing 3 mM biotin by boiling at 95 °C for 10 min under reducing conditions. Proteins were transferred to a nitrocellulose membrane and probed with either a flag tag monoclonal antibody (1:1000; Proteintech, Rosemont, IL) or Alexa Fluor™ 488 streptavidin conjugate (1:3000; Invitrogen, Watham, MA) for detection of biotinylated proteins.

### Single molecule fluorescence in situ hybridization

Brains were rapidly extracted from C57Bl/6J mice and flash-frozen in isopentane on dry ice. Brains were then coronally sectioned at 14 µm thickness in a cryostat. Sections were air-dried for 20 min and subsequently stored at −80 °C until used. The RNAscope Multiplex Fluorescent v2 Assay (Advanced Cell Diagnostics, Newark, CA, USA) was used to visualize mRNA expression following the manufacturer’s instructions except for the washings steps that were extended to 10 min. *Kcnv1* mRNA was detected using the target probe Mm-Kcnv1 (Advanced Cell Diagnostics) and the Opal™ 650 dye (1:1500, Akoya Biosciences, Marlborough, MA, USA)

### Surface Quantification Assay

The surface expression of HA-tagged K_V_7 subunitsin Xenopus oocytes was analysed 2 days after injection of the cRNA (10 ng/oocyte of HA-tagged K_V_7.2 alone or together with 10 ng/oocyte of K_V_8.1 or K_V_8.2). To block non-specific antibody binding, oocytes were incubated in ND96 (in mM: NaCl 96, KCl 2, CaCl2 1.8, MgCl2 1, HEPES 20, Na-pyruvate 2.5, and 100U/ml Penicillin-streptomycin) solution containing 1% Bovine serum albumin (BSA) at 4 °C for 30 min. Oocytes were then incubated for 60 min at 4 °C with 100 μg/ml rat monoclonal anti-HA antibody (clone 3F10, Roche Pharmaceuticals, Basel, Switzerland) in 1% BSA/ND96, washed 6 times at 4 °C with 1% BSA/ND96 and incubated with 2 μg/ml of peroxidase-conjugated, affinity-purified, F(ab)2-fragment goat anti-rat immunoglobulin G antibody (Jackson ImmunoResearch, West Grove, PA) in 1% BSA/ND96 for 60 min. Oocytes were washed thoroughly, first in 1% BSA/ND96 (4°C, 60 min) and then in 1x ND96 without BSA (4 °C, 15 min). Individual oocytes were placed in 20 µ L of SuperSignal Elisa Femto solution (Pierce, Chester, UK) and, after an equilibration period of 10 s, chemiluminescence was quantified in a luminometer (Lumat LB9507, Berthold Technologies, Bad Wildbad, Germany). For each construct, the surface expression of 20 oocytes was analysed in one experiment and at least three experiments (∼40 oocytes) were performed. The luminescence produced by uninjected oocytes was used as a negative control.

### Quantitative reverse transcriptase-PCR

DRG, hippocampus and heart tissue were isolated from C57Bl/6J mice, snap frozen in liquid nitrogen and stored at −80°C until use. RNA extraction and reverse transcription quantitative polymerase chain reaction (RT-qPCR) were performed as previously described (Kalpachidou *et al*, 2022; Kummer *et al*, 2018). Briefly, the peqGOLD TriFast reagent (Peqlab) was used to extract total RNA according to manufacturer’s instructions [chloroform (C2432) and absolute ethanol (107017) were obtained from Merck]. RNA pellets were reconstituted in nuclease free water (R0582, ThermoFisher Scientific) and concentration was measured using NanoDrop 2000 (ThermoFisher Scientific). Reverse transcription was performed using MuLV reverse transcriptase (N8080018, ThermoFisher Scientific) according to the supplier’s protocol. Gene expression was estimated using the following TaqMan Gene Expression Assays (ThermoFisher Scientific): Kcnq1 (Mm00434640_m1), Kcnq2 (Mm00440080_m1), Kcnv1 (Mm00550691_m1), Kcnv2 (Mm00807577_m1), Hprt (Mm00446968_m1), Sdha (Mm01352363_m1), and Tfrc (Mm00441941_m1), with Hprt, Sdha and Tfrc serving as reference genes. Reactions were prepared according to TaqMan Gene Expression Assays protocol (20µ L reactions), loaded on MicroAmp Fast Optical 96-well reaction plates (ThermoFisher Scientific) and run on the 7500 Fast Real-Time PCR System (Thermo Fisher Scientific). The protocol details were as follows: an initial 10 min 95 °C step, followed by 40 two-step cycles of 15 s at 95 °C and 1 min at 60 °C. Samples (eight per tissue) were run as technical duplicates alongside non-template controls. Threshold was set manually at 0.1, whereas baselines were estimated automatically. C_q_ values for each sample were normalized to the respective geometric mean of the C_q_ values of the three reference genes (Hprt/Sdha/Tfrc) and subsequently expressed in relation to the average expression of either Kcnq2 (for DRG and hippocampus tissues) or Kcnq1 (for heart tissue).

### Data analysis and statistics

Patch clamp recordings were analysed with PatchMaster (HEKA) and IgorPro (Wavemetrics, Lake Oswego, OR) or R version 4.1.5 (RCoreTeam, 2014) together with the PatchR package (https://github.com/moritzlindner/PatchR). Voltage dependence of activation was derived from tail current amplitudes using voltage protocols indicated: Maximal amplitudes of tail currents were fit with a two-state Boltzmann function with I = I_min_ + (I_max_-I_min_)/(1 + exp((V-V_h_)/s)), where I is current, V is the membrane voltage, V_h_ is the voltage at half maximal activation, and s describes the steepness of the curve. Results on voltage dependence are shown as normalised tail current (conductance)-voltage curves, obtained by normalizing to (I_max_-I_min_), obtained from fits to data of individual experiments. Statistical analysis was performed with two-tailed Student’s, Wilcoxon signed, Dunnett or Scheffé test as appropriate. For surface expression analysis in oocytes, values from three experiments were normalised to oocytes injected with extracellularly HA-tagged K_V_7 cRNA. Statistical analysis was carried out by ordinary one-way ANOVA with post hoc Dunnett’s multiple comparison test. Statistical significance was assigned at P ≤ 0.05 (* P ≤ 0.05, ** P ≤ 0.01, *** P ≤ 0.001). All data are presented as mean ± SEM. In electrophysiological experiments, *n* represents the number of individual cells recorded from at least three independent experiments (independent transfections or independent tissue, i.e. biological replicates).

Confocal micrographs form PLA experiments were analysed using FIJI (National Institutes of Health, Bethesda, MD, USA, (Schindelin *et al*, 2012)) and a custom-made macro (accessible via: https://www.kvs-liaison.eu/data/PLA_Macro_autothreshold.ijm). This macro enabled unbiased analysis of the micrographs by automatically (1) detecting the boundaries of cells showing flag immunofluorescence, and (2) then counting the PLA dots inside and outside the boundaries of those immunopositive cells separately. Summary statistics were then performed using R version 4.1.5 (RCoreTeam, 2014) and ggplot2 (Wickham H., 2016).

### Bioinformatics

Three individual published single-cell RNA sequencing datasets were utilized to analyse the distribution and co-expression of Kv8, Kv7 and Kv2 in hippocampus (Saunders *et al*., 2018), dorsal root ganglia (DRG, (Finno *et al*., 2019)) and heart (Tabula Muris *et al*., 2018), respectively. Raw gene count matrices were obtained from the Genome Expression Omnibus (GEO) archive (DRG: GSE128276) or author curated archives (hippocampus: dropviz.org, heart: https://figshare.com/projects/Tabula_Muris_Transcriptomic_characterization_of_20_organs_a nd_tissues_from_Mus_musculus_at_single_cell_resolution/27733). Processing and analysis of the datasets were performed using R 4.1.5 and Seurat 3.1 (Usoskin *et al*, 2015). Data were log-normalized, centred, and scaled. Where available, cluster assignments lists as published by the authors of the datasets were utilized. For DRG, clustering was performed using the KNN and Louvain algorithms implemented in Seurat after regressing out technical variance. Obtained clusters were validated using a set of unique cluster markers kindly provided by the authors and the DRG cell-type marker genes as reported by Usoskin et al (Usoskin *et al*., 2015) (Supplementary Figure 6). Dot plots for K_V_7, K_V_2 and K_V_S expression in the individual cell clusters identified in each dataset were drawn with dot radius representing the percentage of cells in an individual cluster expressing the K_V_ or K_V_S gene under investigation and dot colour indicating the mean scaled gene count in that cluster, K_V_/ K_V_S genes detected in less than 10% of the cells belonging to any individual clusters were not visualized. Finally, correlation of the expression levels of K_V_S with K_V_7 and K_V_2 were analysed using Pearson’s correlation coefficients (PCC). Data were bootstrapped (n=100, with cells being individual observations) to obtain an estimate of variance. Data are presented as Box-and-Whisker plots with hinges representing quartiles and whiskers extending to the last data point within the 1.5-fold inter quartile ranges. To explore the biological plausibility of this approach correlation coefficients were also calculated for genes of ion-channels that are known to interact with any of the Kv-channels under investigation (Supplementary Figure 7 D).

## Author Contributions

*Participated in research design: VR, DO, MGL, ML*

*Conducted experiments: VR,* NX, IGS, JS, RB, IS, KK*, MGL, ML*

*Performed data analysis: VR,* NX, IGS, JS, RB, IS, KK*, MGL, ML*

*Wrote or contributed to the writing of the manuscript: VR, KK, MGL, ML*

## Acknowledgements

The authors acknowledge the kind gift of plasmids (K_v_S channels) form Regina Preisig-Müller and Elke Bocksteins and thank Olga Ebers for outstanding technical assistance.

